# Efficacy of transoral laser surgery versus linear accelerator radiotherapy for the treatment of T1 glottic carcinoma: a meta-analysis of oncologic outcomes

**DOI:** 10.1101/2020.04.20.050807

**Authors:** Jiawei Zhu, Jing Shen, Ziye Zheng, Xin Lian, Zheng Miao, Ke Hu, Fuquan Zhang

**Author notes:** These authors contributed equally to this work (Jiawei Zhu, Jing Shen).

## Abstract

**Objective:** A meta-analysis was conducted to compare oncologic outcomes for patients of T1 glottic carcinoma who were treated with transoral laser surgery (TLS) or linear accelerator radiotherapy (linac RT).

**Methods:** All related studies published up to September 2019 were acquired by searching Pubmed, EMBASE, and Cochrane, with the index words: glottic, vocal, laryngeal, radiation, radiotherapy, irradiation, laser, surgery, cordectomy, carcinoma and cancer. Relative studies which compared oncologic outcomes between linac RT and TLS were included. Sensitivity analysis were performed to evaluate heterogeneity.

**Results:** A total of twelve eligible studies were included for the analysis, which contained three prospective studies and nine retrospective studies. Patients who underwent TLS had increased overall survival (OR = 1.40, 95% CI=1.02-1.94, *P*=0.04) and laryngeal preservation (OR = 5.37, 95% CI = 3.05-9.44, *P* < 0.00001) versus who underwent linac RT. No statistical difference was observed between TLS group and linac RT group in terms of local control (OR=0.88, 95% CI = 0.62-1.24, *P* = 0.47), disease-specific survival (OR = 0.61, 95% CI = 0.26-1.43, *P* = 0.26), and disease-free survival (OR = 1.63, 95% CI = 0.70-3.81, *P* = 0.14).

**Conclusions:** The results of this meta-analysis indicate that there were clinical benefits for patients with glottic carcinoma after TLS compared with linac RT with respect to overall survival and laryngeal preservation. However, more multi-center randomized controlled trials would be urgently needed to prove these differences.

## Introduction

Laryngeal carcinoma is the most common malignant tumors of head and neck, with an estimated 211,000 new cases annually [1]. Glottic carcinomas represent two-third of laryngeal carcinoma, which is mainly to the anterior portion of the vocal cord [1]. Most glottic carcinoma can be diagnosed in early stages due to the sparse lymphatics and the early symptoms affecting vocal cord such as persistent hoarseness.

The primary aim of treatment for glottic carcinoma is to achieve local control and increase survival, while the secondary aim is to preserve the organ as well as its functions. Transoral laser surgery (TLS) and linear accelerator radiotherapy (linac RT) are the two main effective treatment modalities for early glottic carcinoma with high cure rates [2–4]. There is still a lack of standard guiding therapeutic selection in early glottic carcinoma, adding to the complexity of decision making. Previous meta-analyses suggested that TLS was significantly better in laryngeal preservation and overall survival for T1 glottic carcinoma, while RT resulted in better voice quality [5–7]. The results on disease-specific survival and disease-free survival were conflict [5, 7]. In addition, most meta-analysis reported heterogeneity on the outcomes of local control and laryngeal preservation [4, 5, 7, 8]. Therefore, we perform this study with restrict inclusion criteria and updated literature comparing only TLS and linac RT, which are now the major treatment modality clinically.

## Method

### Data source, search strategy, and selection criteria

This study was conducted and reported according to the Preferred Reporting Items for Systematic Reviews and Meta-Analysis Statement issued in 2009 (Checklist S1) [9]. Studies that compared the outcomes between TLS and linac RT for the treatment of T1 glottic carcinoma were selected for meta-analysis in this study. There is no language or publication status limitation for literature review. A systematic electronic search of PubMed, EmBase, and the Cochrane Library databases was performed for eligible studies from inception to September 2019. The following index words were used: glottic, vocal, laryngeal, radiation, radiotherapy, irradiation, laser, surgery, cordectomy, carcinoma and cancer. We also reviewed the reference lists of the included studies for undetected relevant studies.

The retrieved records were independently reviewed by two investigators. If the investigators disagreed about the eligibility of an article, it was resolved by debating with a third reviewer. The inclusion criteria were as follows: (1) patients with T1 glottic cancer; (2) intervention was TLS compared with linac RT; (3) outcomes data reported; and (4) article type as original research. In the current study, the outcome included tumor outcomes, such as local control, laryngeal preservation, overall survival, disease-specific survival and disease-free survival.

### Data collection and quality assessment

Research articles were evaluated by two independent researchers, and any inconsistencies were discussed before reaching a consensus. For each study, the following data was collected: the first author’s name, publication year, country where the study was conducted, study design, number of patients, age, gender, tumor stage, treatment, and follow-up period as the baseline data. Primary outcomes were oncological outcomes including local control, laryngeal preservation, overall survival, disease-specific survival and disease-free survival.

The Newcastle-Ottawa Scale (NOS) was used to evaluate the quality of the included studies for cohort studies. The NOS was based on the following three subscales: selection (four items), comparability (one item), and outcome (three items). NOS ranges were 0 to 8, and scores of 0–3, 4–6, and 7–8 were considered as low, moderate, and high quality, respectively, for these studies.

### Statistical analysis

The results from each study were considered as dichotomous frequency data, and odds ratio (OR) and 95% confidence intervals (CIs) of each individual trial were calculated from event numbers in each group within the individual trials before data pooling. Furthermore, ORs and 95% CIs were calculated for overall survival, laryngeal preservation, local control, disease-specific survival, and disease-free survival in glottic cancer patients receiving TLS and linac RT. The potential heterogeneity across the studies was examined using the Cochran’s *Q*-statistic and I^*2*^ statistics. If the *P* value for heterogeneity was < 0.1 or I^2^ > 50%, then it was considered as significant heterogeneity. A fixed effect model was used to combine all the study results. Sensitivity analyses were performed to explore the sources of heterogeneity of the included studies by removing each included study subsequently. Publication bias was evaluated using the Egger’s and Begg’s test [10], and *P* < 0.05 was considered as a statistically significant publication bias; funnel plots were also conducted [11]. All reported *P* values were two-sided, and *P* < 0.05 were considered significant for the included studies. Statistical analyses were performed with STATA software (version 15.0; Stata Corporation, College Station, TX, USA).

## Result

### Characteristics of the included studies

A total of 1223 studies were retrieved based on our searching strategy. After reading the abstract, 27 studies were related to our aims (Fig.1), and 15 of them were subsequently excluded with reasons give. The remaining 12 studies were selected for our analysis (Table 1) [12–23]. A total of 1496 participants were selected for the meta-analysis, of which 753 and 743 patients underwent TLS and linac RT, respectively. None of the 12 studies included were completely randomized, 3 were prospective studies, and the remaining were retrospective. The majority of the studies were from Europe and North America. All patients were diagnosed with T1 (T1a, T1b) glottic carcinoma.

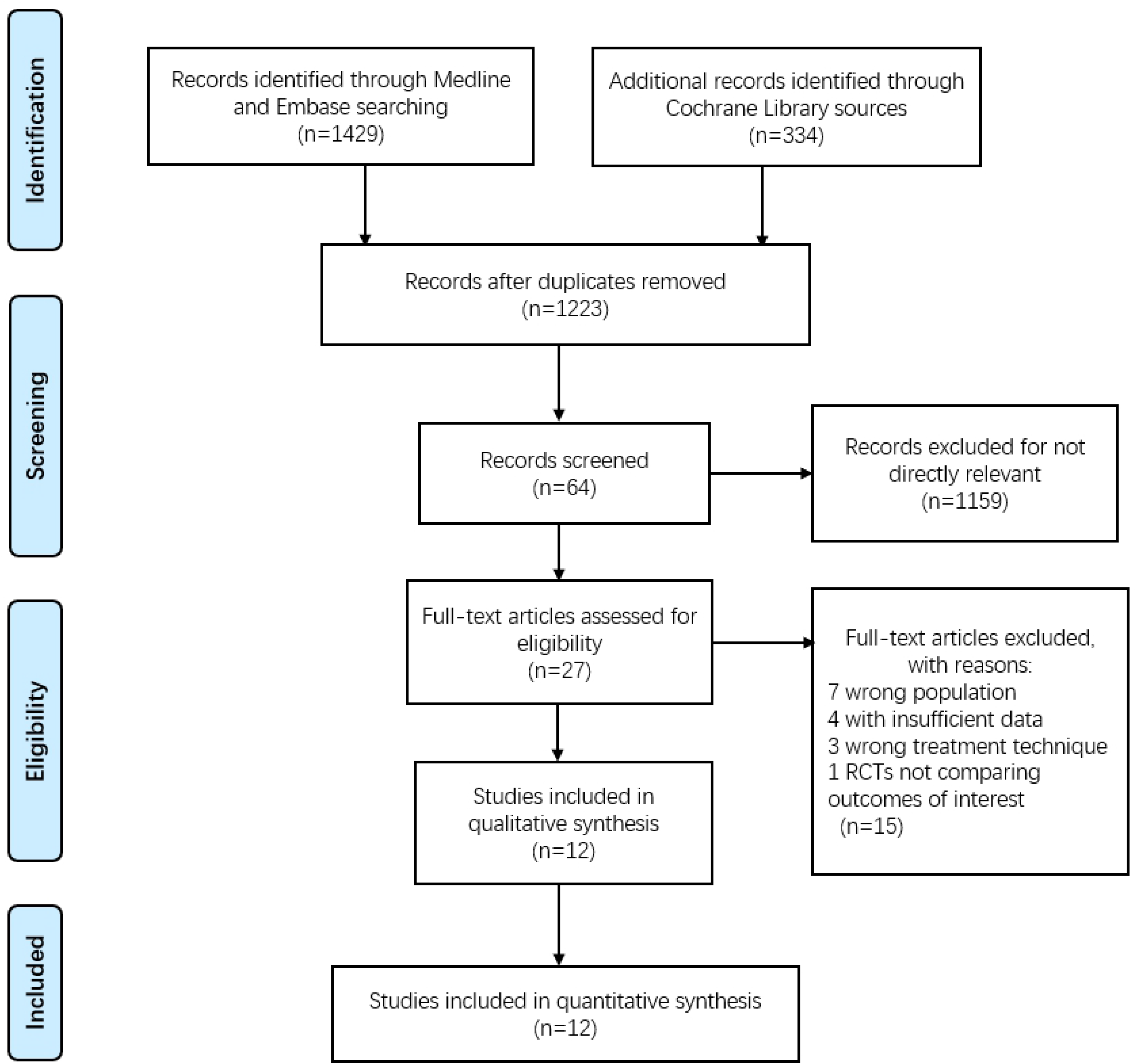

**Table 1.**
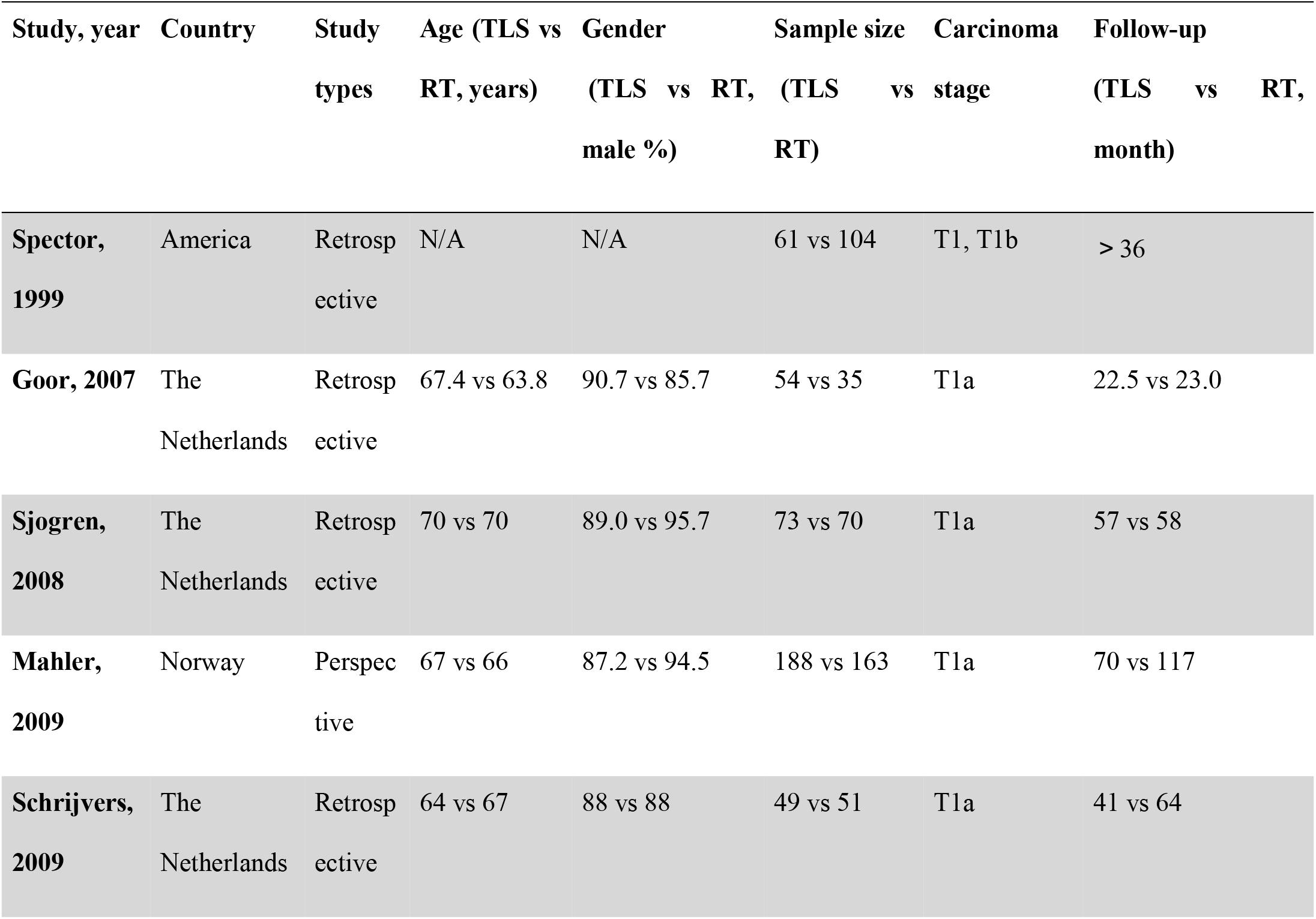
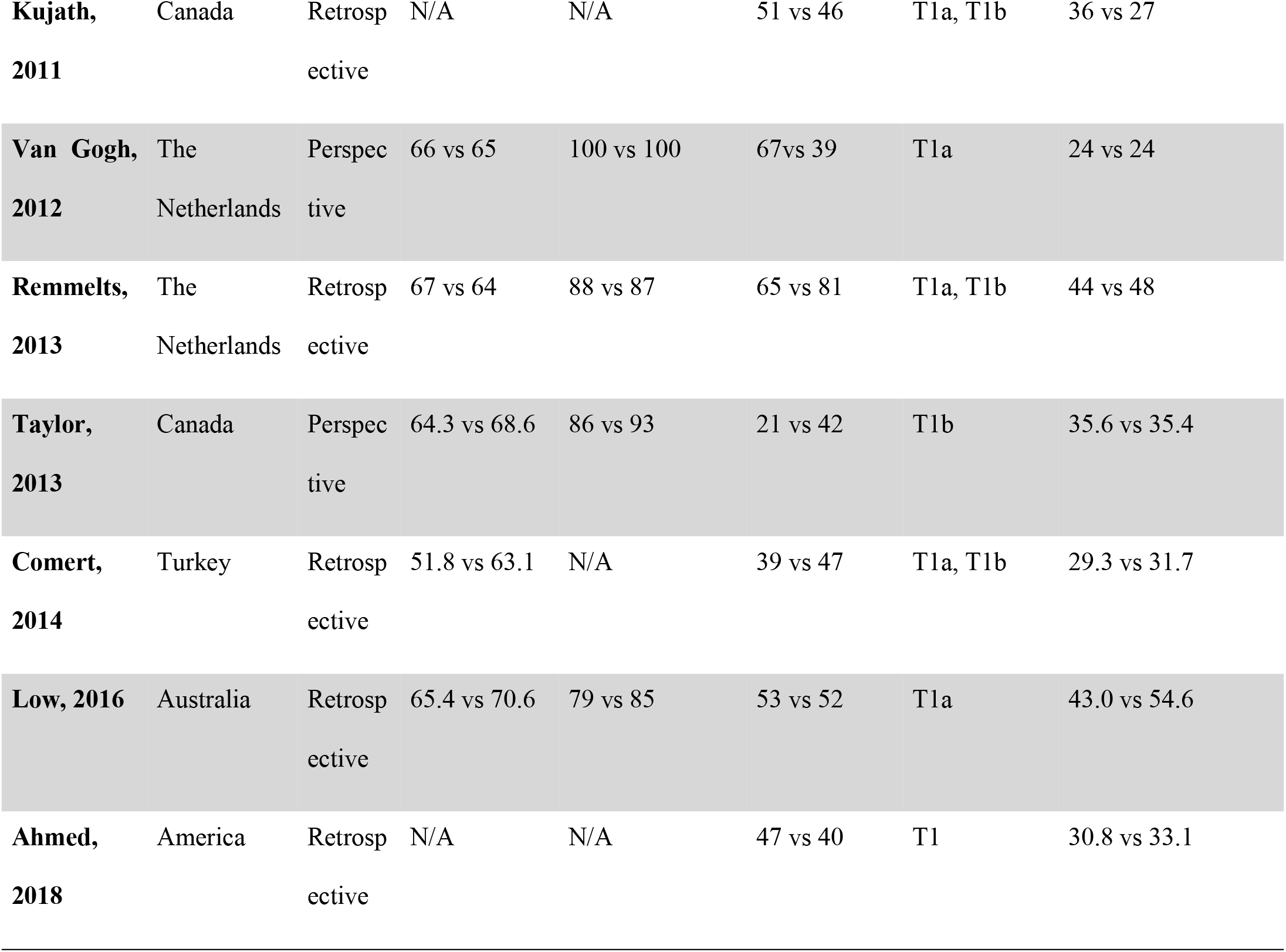
Characteristics of the included studies.

NOS was used to examine the quality of all included studies. Half of the studies were marked as 7 in NOS as they failed to report the adequacy of follow-up, and 2 studies [17, 20] were marked as 6 due to the inadequacy of follow-up and the lack of detailed treatment method. The 4 remaining studies received full scores in NOS, indicating they were high quality original studies (Table 2).

**Table 2.**
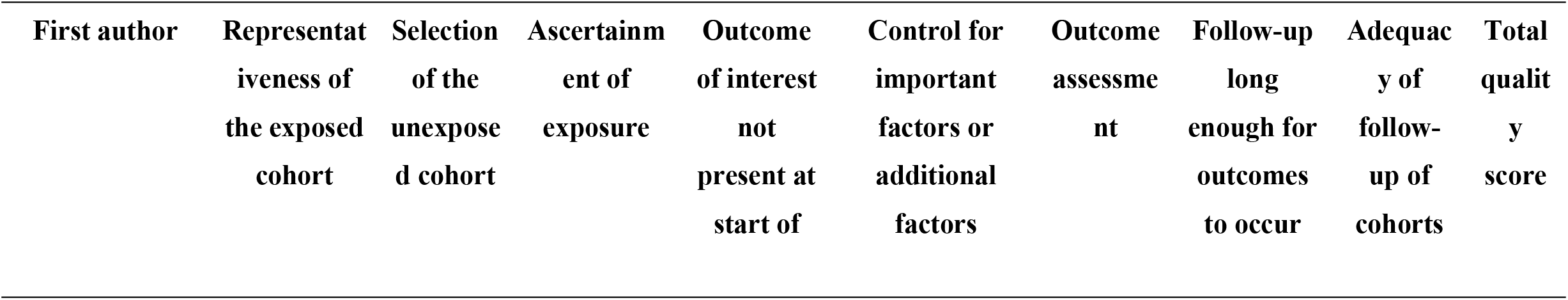
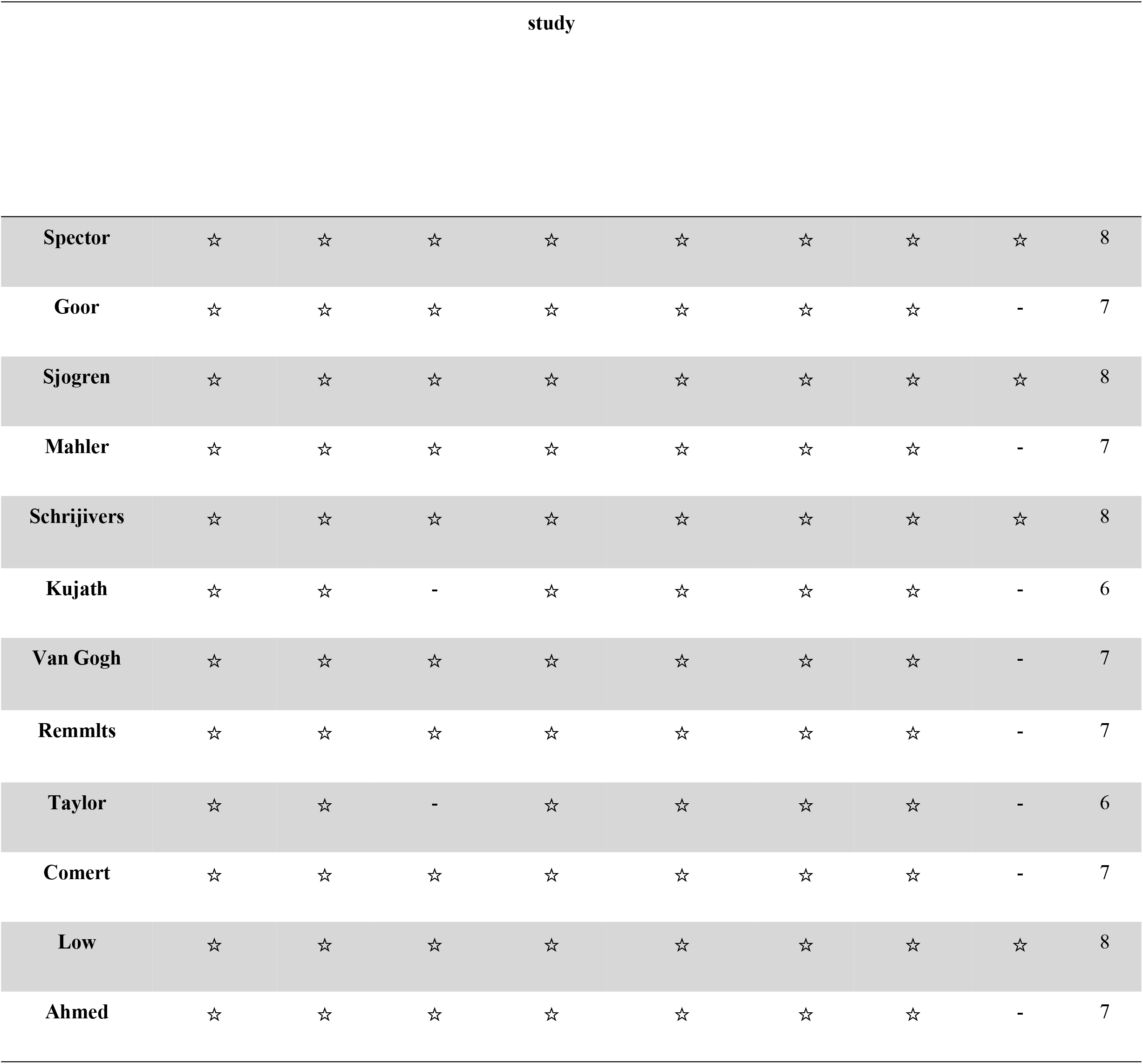
Methodological quality of the studies included in the meta-analysis.

| First author | Representativeness of the exposed cohort | Selection of the unexposed cohort | Ascertainment of exposure | Outcome of interest not present at start of | Control for important factors or additional factors | Outcome assessment | Follow-up long enough for outcomes to occur | Adequacy of follow-up of cohorts | Total quality score |
| --- | --- | --- | --- | --- | --- | --- | --- | --- | --- |

| study |  |  |  |  |  |  |  |  |  |
| --- | --- | --- | --- | --- | --- | --- | --- | --- | --- |
| Spector | ☆ | ☆ | ☆ | ☆ | ☆ | ☆ | ☆ | ☆ | 8 |
| Goor | ☆ | ☆ | ☆ | ☆ | ☆ | ☆ | ☆ | - | 7 |
| Sjogren | ☆ | ☆ | ☆ | ☆ | ☆ | ☆ | ☆ | ☆ | 8 |
| Mahler | ☆ | ☆ | ☆ | ☆ | ☆ | ☆ | ☆ | - | 7 |
| Schrijvers | ☆ | ☆ | ☆ | ☆ | ☆ | ☆ | ☆ | ☆ | 8 |
| Kujath | ☆ | ☆ | - | ☆ | ☆ | ☆ | ☆ | - | 6 |
| Van Gogh | ☆ | ☆ | ☆ | ☆ | ☆ | ☆ | ☆ | - | 7 |
| Remmlts | ☆ | ☆ | ☆ | ☆ | ☆ | ☆ | ☆ | - | 7 |
| Taylor | ☆ | ☆ | - | ☆ | ☆ | ☆ | ☆ | - | 6 |
| Comert | ☆ | ☆ | ☆ | ☆ | ☆ | ☆ | ☆ | - | 7 |
| Low | ☆ | ☆ | ☆ | ☆ | ☆ | ☆ | ☆ | ☆ | 8 |
| Ahmed | ☆ | ☆ | ☆ | ☆ | ☆ | ☆ | ☆ | - | 7 |

### Overall survival

Of the twelve studies included in our meta-analysis, nine studies provided detailed data on overall survival with 596 patients in the TLS group and 611 patients in the RT group. Meta-analysis revealed low heterogeneity among the eight retrospective and one prospective cohort studies (I^2^ = 0%, *P* = 0.95), and the fixed effect model was applied. Pooled analysis demonstrated that participants had a favorable overall survival in the TLS group in comparison with the RT group (OR = 1.40, 95% CI = 1.02-1.94, *P* = 0.04; Fig. 2).

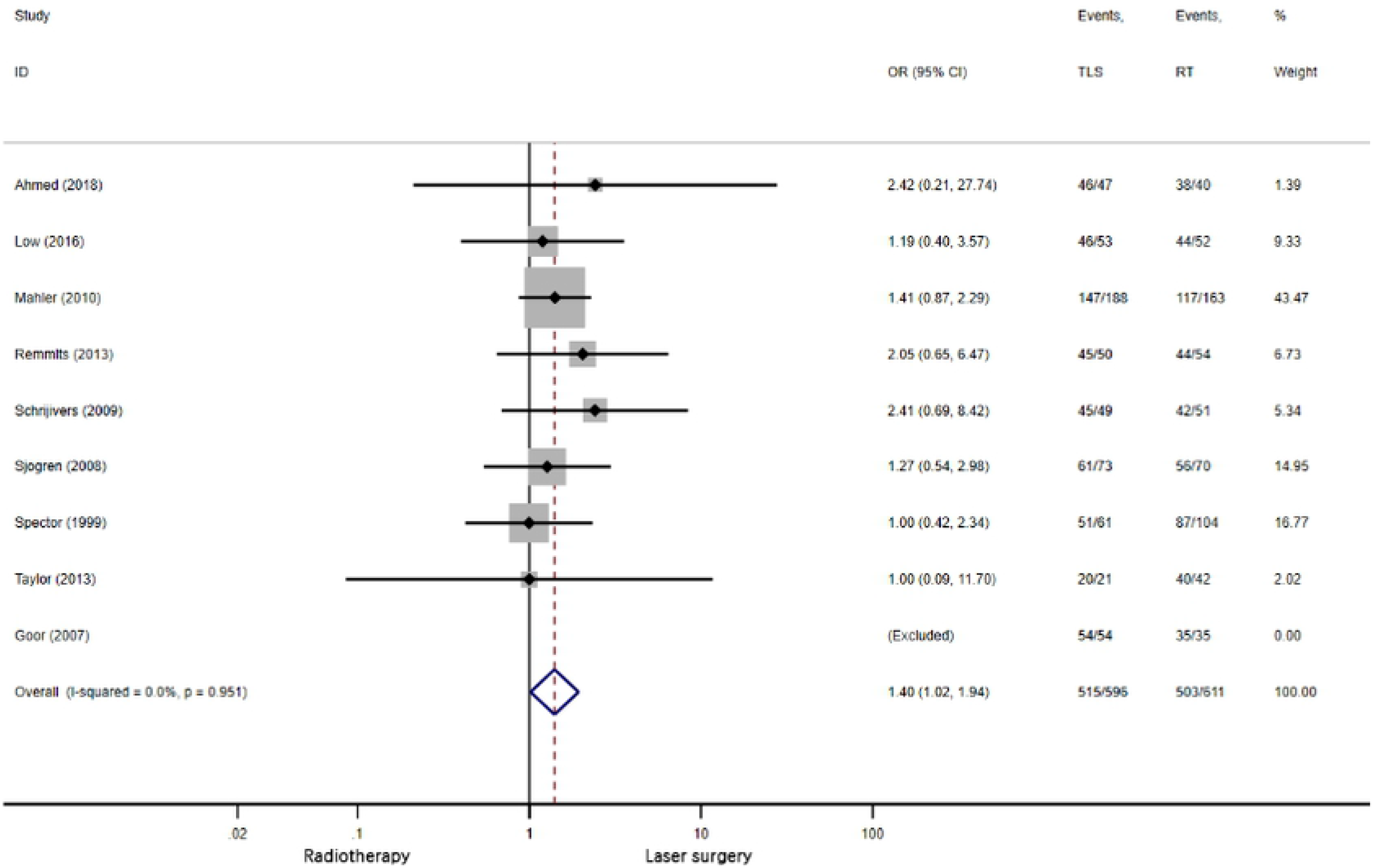

### Larynx preservation

A total of eleven studies having 714 patients in the TLS group and 692 patients in the RT group were evaluated for laryngeal preservation. There was low heterogeneity among the eight retrospective and three prospective studies (I^2^ = 17%, *P* = 0.28), and the fixed effect model was applied. This meta-analysis showed a strong favorable outcome for laser treatment in terms of laryngeal preservation (OR = 5.37, 95% CI = 3.05-9.44, *P* < 0.00001; Fig. 3).

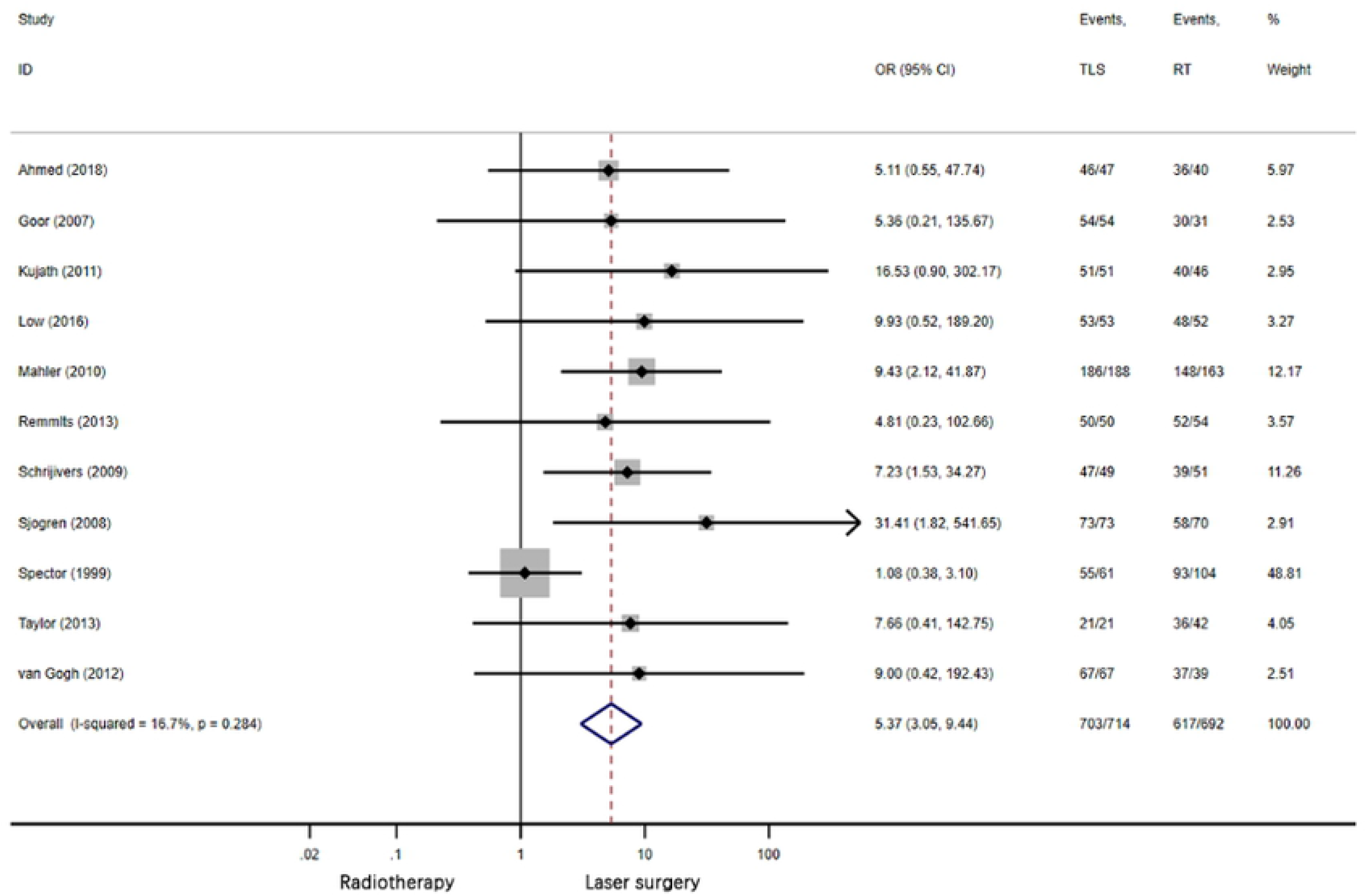

### Local control

Eleven studies having 693 patients in RT group and 700 patients in the LS group reported local control. There was low heterogeneity among the nine retrospective and two prospective studies (I^2^=26%, *P* = 0.19), and the fixed effect model was applied. Results of the pooled effect demonstrated that the difference between RT and TLS on local control was not statistically significant (OR = 0.88, 95% CI = 0.62-1.24, *P* = 0.47; Fig. 4).

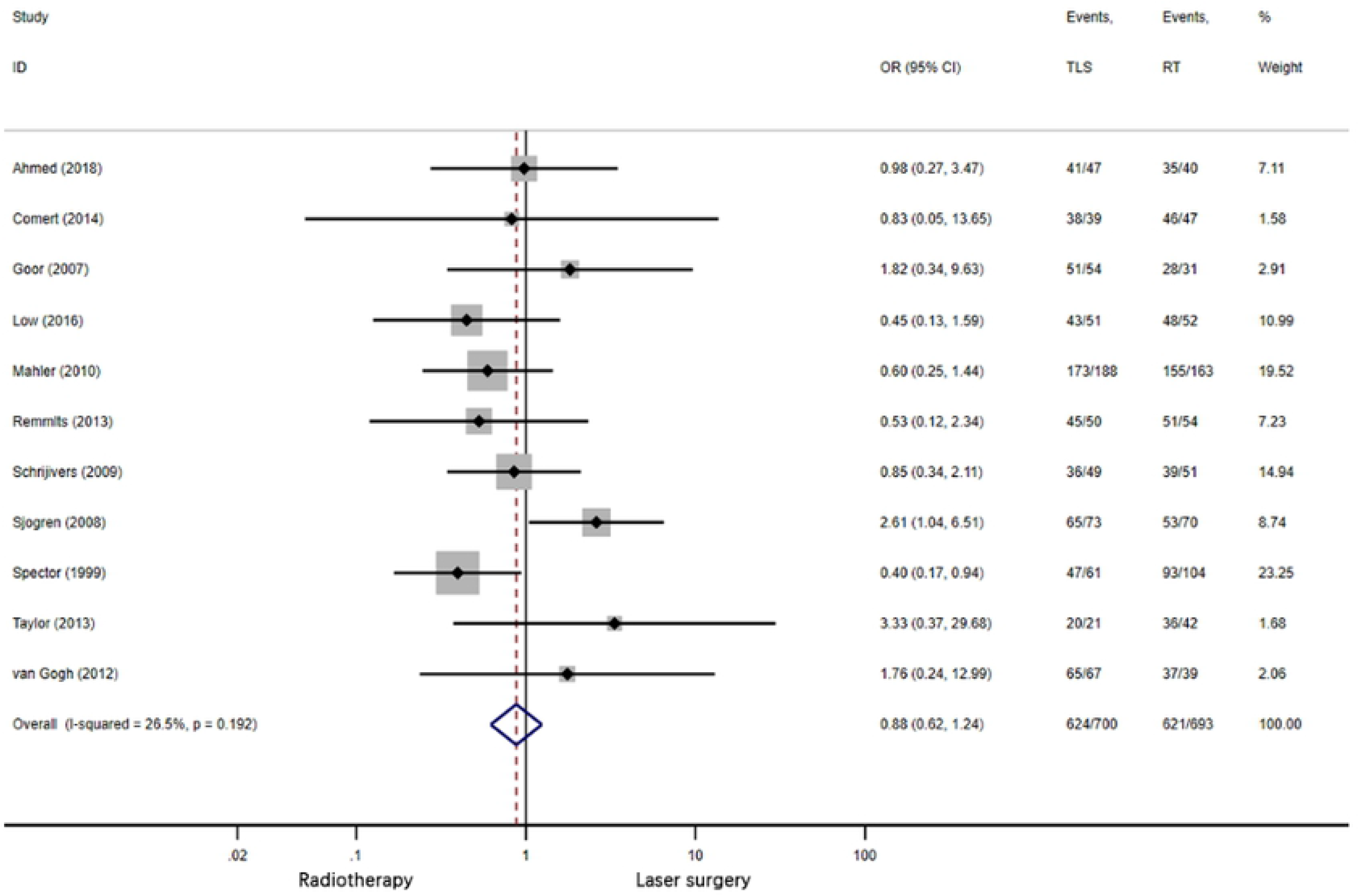

### Disease-specific survival

Seven of twelve studies reported event numbers for disease specific survival with 459 patients in the RT group and 455 patients in the TLS group. There was insignificant heterogeneity was observed (I^2^ = 0%, *P* = 0.86), and no statistical significance (OR = 0.61, 95% CI = 0.26-1.43, *P* = 0.26; Fig. 5).

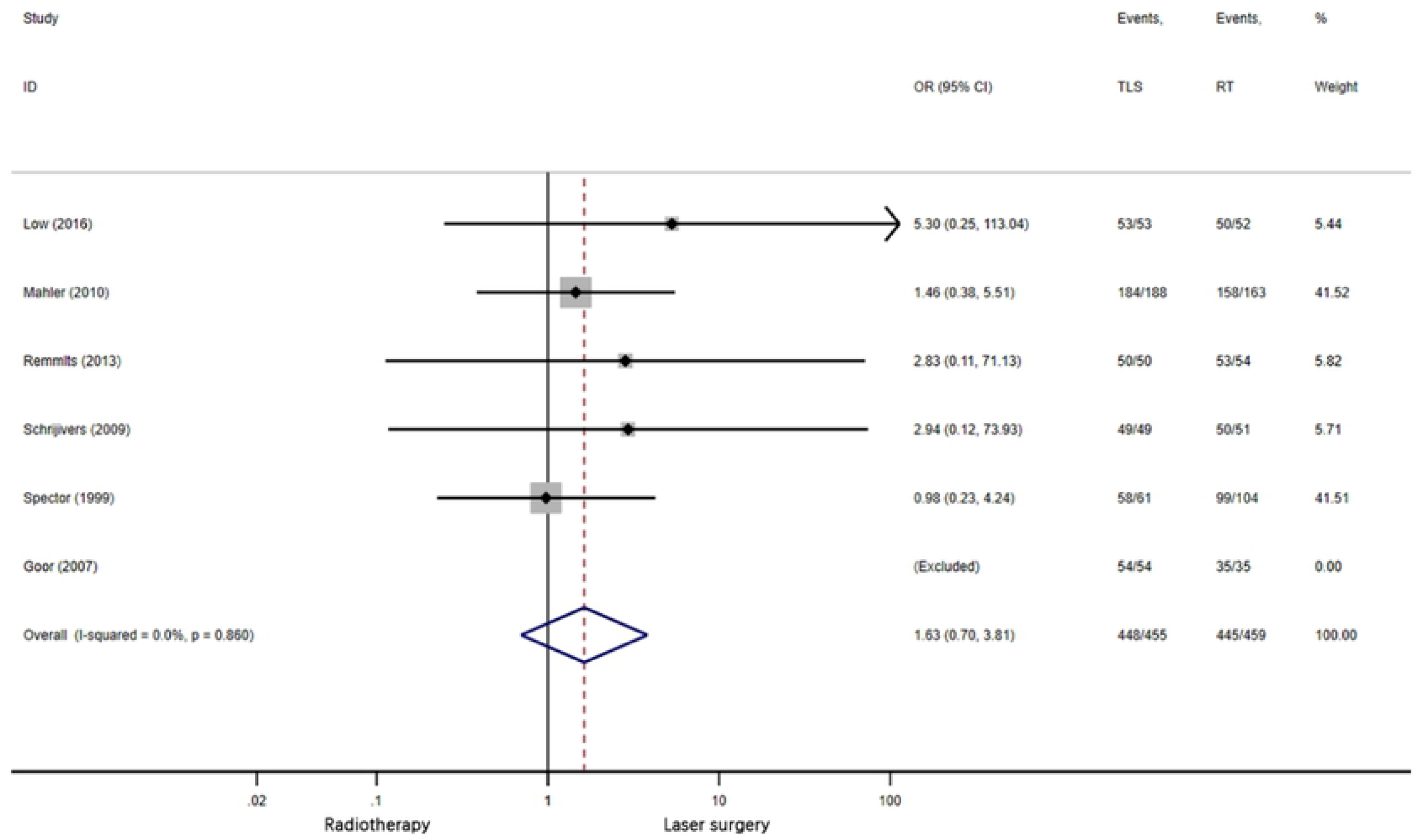

### Disease-free survival

Five studies having 344 patients in RT group and 348 patients in the LS group reported disease-free survival. There was low heterogeneity among the five retrospective studies (I^2^ = 8%, *P* = 0.36), and the fixed effect model was applied. Results of the pooled effect demonstrated that the difference between RT and LS on disease-free survival was not statistically significant (OR = 1.63, 95% CI = 0.70-3.81, *P* = 0.14; Fig. 6).

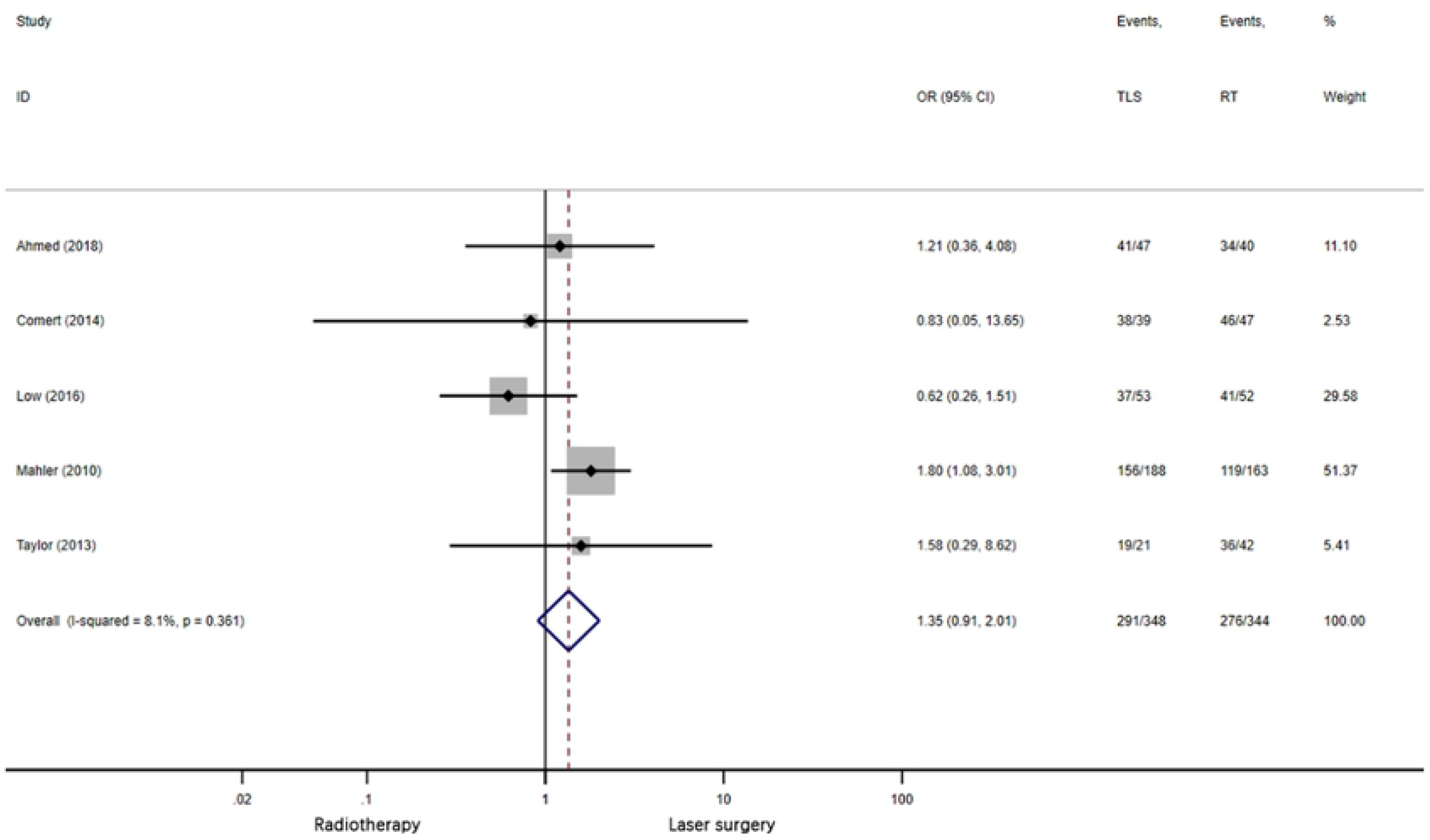

### Sensitivity analysis

Sensitivity analysis was performed using the leave-one-out approach, in which the analysis was performed repeatedly with each removed once (Supplementary Fig. 2). The direction and magnitude of combined estimates for laryngeal preservation, local control and disease-specific survival did not vary markedly with the removal of any one study, indicating that these findings were robust and the data were not overly influenced by any given study. For overall survival, removal of four studies [15, 16, 19, 23] resulted in the difference between treatments becoming insignificant; however, removal of all other studies resulted in no change in magnitude or direction the overall survival. For disease-free survival, removal of Low et al [22] resulted in the difference between treatments becoming significant (*P* = 0.03); removal of Mahler et al [15] resulted in the change in magnitude of the disease-free survival. Overall, the sensitivity analysis indicated that no one study overly influenced the pooled estimates, demonstrating that the findings are robust.

In addition, the two studies without detail treatment technique provided result on larynx preservation at the same time. After excluding them, the result for larynx preservation was not affected. (supplementary Fig. 3).

### Publication bias

Review of the funnel plots could not rule out the potential for publication bias (Supplementary fig. 4). The Begg’s test showed no evidence for publication bias. However, Egger’s test showed laryngeal preservation had potential publication bias (*P* = 0.021). A sensitivity analysis was conducted with trim and fill method, and the result was the same [24].

## Discussion

Although laryngeal carcinoma has been high prevalence, most patients can be diagnosed at an early stage and rarely presents metastasis [5]. TLS and RT are both effective methods for treating early glottic carcinoma with comparable cure rates [25]. Each treatment modality has its advantages and disadvantages.

TLS allows targeted resection with cheaper expense, shorter period and the histological confirmation of surgical margins [26]. It can also be repeated as a salvage therapeutic for recurrence [27, 28]. The disadvantages of TLS include: the need of special surgical techniques, exposure of endolarynx, removal of extra normal tissue with oncologic margin, and surgical anesthetic risks. At present, CO2 laser is the mainstay of TLS, while potassium titanyl phosphate (KTP) laser has become an emerging treatment modality. Several studies using KTP laser have shown similar oncological outcomes for glottic carcinoma as CO2 laser and RT [23, 30, 31], and better subjective vocal outcomes than CO2 laser [30]. As our study focused on the oncological outcomes, we did not limit the type of laser. Two retrospective cohorts implementing KTP and one with microscopic diode laser surgery (MDLS) were included.

Compared with TLS, RT may in better voice quality due to the more intact preservation of vocal cords [29] without the restriction of laryngeal exposure and general anesthesia. However, RT cannot be repeated after recurrence, and the salvage is a partial or total laryngectomy. In addition, the entire larynx is irradiated, causing actinic edema [5]. In terms of radiation techniques, Co-60 teletherapy has historically been implemented, but it fell out favor at the end of 20^th^ century because of the prevalent use of linac RT [32], which provides a greater penetration, higher dose rate and better radiation safety [33]. Different types, segmentation mode and radiation dose of RT were included in previous systematic reviews. Researches comparing linac RT with TLS were included, and studies with Co-60 were excluded for more reliable results.

The lack of prospective randomized controlled trial comparing TLS and RT adds to the complexity of therapy selection. Previous systematic reviews with meta-analyses showed that TLS was favourable for OS and laryngeal preservation [4–7]. However, all of them revealed heterogeneities in oncologic outcomes. By dividing the studies into subgroups based on publication year of 2000, Huang et al [4]. found no heterogeneity and contrary results on laryngeal preservation in each subgroup. This indicated that with the development of modern laser and radiation techniques, different treatment modalities might result in the heterogeneity partially.

Our meta-analysis revealed the absence of heterogeneity in all oncological outcomes. The results confirmed that TLS was equally efficacious as linac RT in local control, disease-specific survival and disease-free survival. TLS is favorable for laryngeal preservation and overall survival. This conclusion was unchanged after sensitive analysis and subgroup analysis. Patients treated with TLS are therefore approximately five times more likely to preserve their larynx than those treated with linac RT. Several previous systematic reviews with meta-analysis have reported similar results, which suggested that TLS was superior to preserve the larynx for patient with glottic carcinoma [4, 6–8]. This advantage could arise from the ability to precisely resect lesions and conserve surrounding anatomy with TLS. In addition, the repeatability of TLS for recurrence also contribute to its higher laryngeal preservation [7, 23]. In terms of overall survival, three systematic reviews supported that TLS was advantageous when compared with RT, which might be associated with the treatments after recurrence. Single-arm retrospective studies also presented preferable outcomes in patients treated with primary TLS than patients with primary RT [34, 35].

The quality of the present meta-analysis is limited by the quality of available literature. First, all included studies were non-randomized cohort studies with only two prospective studies. There might be selection bias due to the physician’s or patient’s preference. Tumors treatable by TLS are relatively more superficial and localized, and physicians are inclined to select these patients into studies [18]. Second, the included studies covered a long period with a relatively small number of patients. Diagnosis, health care and treatment management regarding glottic carcinoma have advanced over time, leading to heterogeneity between studies. Third, the inconsistence in the detail of TLS and RT method between different studies also increased the heterogeneity.

In summary, T1 stage glottic carcinoma patients treated with TLS had increased overall survival and laryngeal preservation compared to patients treated with linac RT. There was no difference in local control, disease-specific survival, and disease-free survival between TLS and linac RT. The ability to repeat TLS or radiation after initial TLS might explain TLS to be a better option with improved overall survival and laryngeal preservation. Taking the surgical contraindications and voice function into account, RT is still an important treatment modality for glottic carcinoma. Therefore, prospective large-scale randomized controlled trial is required to address the remaining uncertainties and establish clinical guidelines.

## Supporting information

S1 File. PRISMA checklist.

(PDF)

S2 File. Sensitive analysis.

(TIF)

S3 File. Forest plot of laryngeal preservation after excluding two studies without treatment details.

(TIF)

S4 File. Funnel plots.

(TIF)

S5 File. S5 Begg’s and Egger’s test & trim and fill method for laryngeal preservation.

(TIF)

## Author Contributions

### Conceptualization

Jiawei Zhu, Jing Shen, Ziye Zheng, Zheng Miao, Xin Lian, Ke Hu, Fuquan Zhang

### Data curation

Jiawei Zhu, Jing Shen, Ziye Zheng

### Formal analysis

Jiawei Zhu, Ziye Zheng

### Methodology

Jiawei Zhu, Jing Shen, Zheng Miao, Xin Lian, Fuquan Zhang

### Project administration

Jiawei Zhu, Jing Shen

### Supervision

Ke Hu.

### Visualization

Jing Shen, Ziye Zheng, Zheng Miao, Xin Lian, Fuquan Zhang

### Writing – original draft

Jiawei Zhu, Ziye Zheng

### Writing – review & editing

Jiawei Zhu, Jing Shen, Zheng Miao, Xin Lian, Ke Hu, Fuquan Zhang

